# Multiscale 3-dimensional pathology findings of COVID-19 diseased lung using high-resolution cleared tissue microscopy

**DOI:** 10.1101/2020.04.11.037473

**Authors:** Guang Li, Sharon E. Fox, Brian Summa, Bihe Hu, Carola Wenk, Aibek Akmatbekov, Jack L. Harbert, Richard S. Vander Heide, J. Quincy Brown

## Abstract

The study of pulmonary samples from individuals who have died as a direct result of COVID-19 infection is vital to our understanding of the pathogenesis of this disease. Histopathologic studies of lung tissue from autopsy of patients with COVID-19 specific mortality are only just emerging. All existing reports have relied on traditional 2-dimensional slide-based histological methods for specimen preparation. However, emerging methods for high-resolution, massively multiscale imaging of tissue microstructure using fluorescence labeling and tissue clearing methods enable the acquisition of tissue histology in 3-dimensions, that could open new insights into the nature of SARS-Cov-2 infection and COVID-19 disease processes. In this article, we present the first 3-dimensional images of lung autopsy tissues taken from a COVID-19 patient, including 3D “virtual histology” of cubic-millimeter volumes of the diseased lung, providing unique insights into disease processes contributing to mortality that could inform frontline treatment decisions.

## Introduction

Coronavirus disease (COVID-19) is a respiratory illness caused by the novel coronavirus SARS-CoV-2, which has sparked a global pandemic with global cases and deaths as of May 5, 2020 reaching to over 3,680,000 and 258,000, respectively, with both numbers rapidly climbing. At the time of this writing, in the US there have been over 72,000 deaths attributed to COVID-19. In severe cases, complications of the disease include severe pneumonia, respiratory failure, acute respiratory distress syndrome (ARDS), and cardiac injury.^1^ There have been limited reports of pathological findings at autopsy from patients who have died as a result of COVID-19. An early case report from China from a single patient analyzed post-mortem biopsies taken from the lung, liver, and heart of the patient.^2^ Analysis of pulmonary biopsy H&E histology indicated diffuse alveolar damage and hyaline membrane formation indicative of ARDS, as well as interstitial lymphocyte infiltration and the presence of multinucleated syncytial cells in the intra-alveolar spaces showing viral cytopathic-like changes. More recently, a report from one of our institutions (LSU Health Sciences Center - University Medical Center), representing the first pathologic analysis of dissected lung and cardiac tissues from open autopsy, confirmed that in addition to diffuse alveolar damage, a novel finding of pulmonary thrombotic microangiopathy and hemorrhage significantly contributed to death, with evidence of the activation of megakaryocytes contributing to small vessel clot formation.^3^

While analysis of sections on slides remains the gold-standard for histological examination, limitations of 2-dimensional sectioning may create artifacts that complicate interpretation. Fully 3-dimensional analysis of the microstructure of tissue samples can provide helpful disambiguation of such sectioning artifacts, and may further positively augment pathologic interpretation.^4^ Three-dimensional histology can be obtained by exhaustive serial sectioning of a tissue block, followed by digital whole slide scanning of the sections and image reconstruction, but such methods are prohibitively expensive and time consuming. Emerging techniques using fluorescence contrast to simulate H&E staining,^5,6^ optical tissue clearing,^4,7–9^ and high throughput optical sectioning microscopes,^4,5,7–11^ have enabled acquisition of 3-dimensional pseudo-histological imaging of a wide range of tissues. In this work, we report the first 3-dimensional microscopy images from optically-cleared lung tissues taken from autopsy of patients who have died as a result of COVID-19 infection. Consent for autopsy was non-restricted by the next of kin, and these studies on autopsy tissues were determined to be exempt by the IRB at LSU Health Sciences Center and Tulane University. We demonstrate near-teravoxel volumetric images of optically-cleared lung tissue over cubic millimeter volumes with virtual H&E contrast and at submicron resolution. Our work builds on the prior autopsy case-series report from LSU-HSC, confirming and expanding novel findings of widespread pulmonary microangiopathy involving activation of mature megakaryocytes in the lung.

## Results and Discussion

**Figure 1** contains reconstructed volumes, selected optical slices, and 3D renderings from a 7.8 mm × 5.9 mm × 0.9 mm formalin-fixed block of lung tissue sampled from the left peripheral upper lobe. Tissue blocks were stained with TO-PRO-3 for nuclear contrast and Eosin-Y for cytoplasmic/stromal contrast, followed by optical clearing with ethyl-cinnamate, according to the method of Reder et al.^7^ Blocks were then imaged using one side of a dual-inverted selective plane illumination microscope^12^ customized for large cleared tissue imaging, described previously.^8,11^ Two grayscale images were collected corresponding to each fluorescent label, which were reconstructed independently, and synthesized into a single pseudo-colored RGB image replicating the color palette and appearance of H&E staining using the approach described by Giacomelli et al. (**Supplemental Figure 1**).^5^ The total raw volume in **Figure 1a** comprised 929 gigavoxels, with a voxel size of 0.363 μm × 0.363 μm × 0.363 μm. Visualization of the entire volume enables distinction of features of multiple scales, from large branching vessels of millimeter dimension, down to single cell nuclei (**Supplemental Figure 2**). A smaller volume (1.2 mm × 0.6 mm × 0.3 mm comprising 4.5 gigavoxels) was selected from the larger volume for detailed visualization. **Supplemental Video 1** shows all slices from all orthogonal orientations in the sub-volume. **Figure 1b** shows a single XY slice from the smaller sub-volume - the location of the slice in the volume is shown in the inset of Figure 1a. Prominent features observed in this virtual H&E slice included a megakaryocyte in a branching small vessel, apparent multi-nucleated or grouped cells demonstrating viral cytopathic changes including prominent nucleoli in the alveolar space, and scattered hyaline-fibrin aggregates in the alveolar spaces. Further analysis of these areas in 3D (**Figure 1c**) provided additional value in interpretation, confirming that the megakaryocyte displayed the expected lobular, fused multi-nuclear shape (**Supplemental Video 2**), enabling determination that the apparent multi-nucleated cell with viral cytopathic effect in Figure 1b was in fact a cluster of independent cells, and showing a spherical hyaline-fibrin aggregate surrounded by apparent inflammatory cells in the alveolar space, consistent with diffuse alveolar damage.

**Figure 1.**
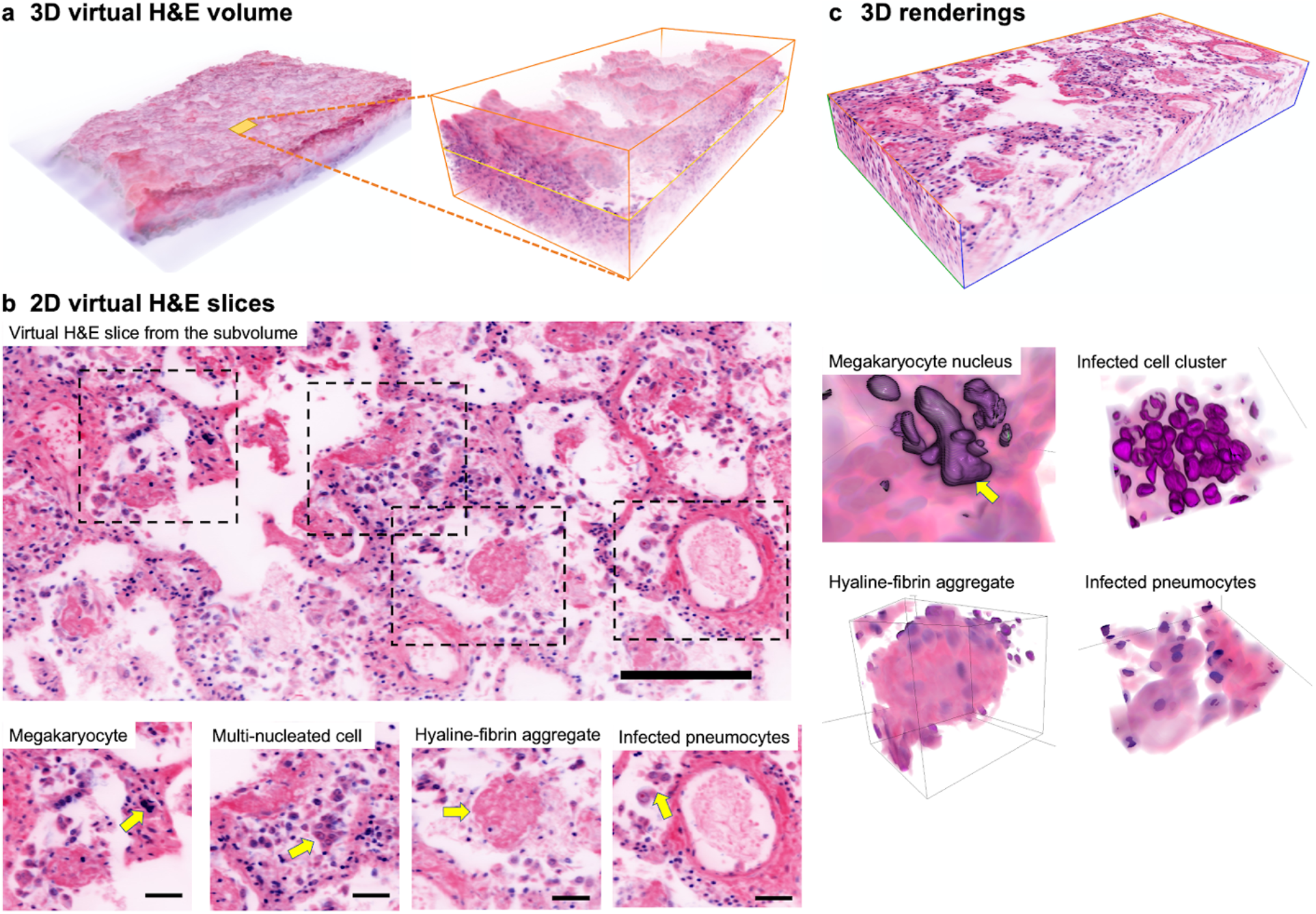
Multiscale 3-dimensional imaging of lung tissue demonstrating viral cytopathic changes, diffuse alveolar damage, and microangiopathy. a) Full 3-dimensional virtual H&E volume of a single gross slice of optically-cleared and fluorescently-stained lung tissue, measuring 7.8 mm × 5.9 mm × 0.9 mm and comprising 0.929 teravoxels of raw image data. A smaller volume (1.2 mm × 0.6 mm × 0.3 mm comprising 4.5 gigavoxels) was selected for detailed analysis (right inset). b) 2-dimensional virtual H&E sections of the smaller volume from (a). Megakaryocyte in the capillaries, infected cell clusters and infected pneumocytes in the alveolar spaces, and hyaline-fibrin aggregates in the alveoli are apparent (yellow arrows). c) Selected 3-dimensional renderings corresponding to structures identified by yellow arrows in (b). 3D renderings of the tissue reveal nuclear, cellular, and tissue morphology in 3D.

An important finding from the initial case-series of autopsies of African-American patients with COVID-19-specific mortality was the presence of pulmonary microangiopathy, apparently involving activated megakaryocytes in the lung.^3^ **Figure 2** contains 3D renderings and optical slices taken from a different subvolume of the tissue, this one measuring 1.65 mm × 0.6 mm × 0.3 mm and comprising 6 gigavoxels. **Figure 2a** shows a 3D rendering of the volume, and **Figure 2b** shows orthogonal slices sampled from the volume as indicated in Figure 1a. This area of the tissue is remarkable for extensive platelet-rich clotting with adherent mononuclear cells in a branching vessel, supplying capillaries surrounding alveoli (yellow arrows). As seen in the orthoslices, the clot can be seen extending throughout the entire length of the vessel sampled in this volume. In addition, extensive fibrin clotting in the small capillaries surrounding the alveoli are observed (green arrows). **Supplemental Video 3** shows orthoslices through the entire volume at each orientation, where the extent of the clotting in the vascular network is clearly observed. Additionally, **Figure 2c** shows a volume render of the tissue, in which the extent of the platelet-rich clotting with adherent inflammatory cells in this subvolume may be appreciated.

**Figure 2.**
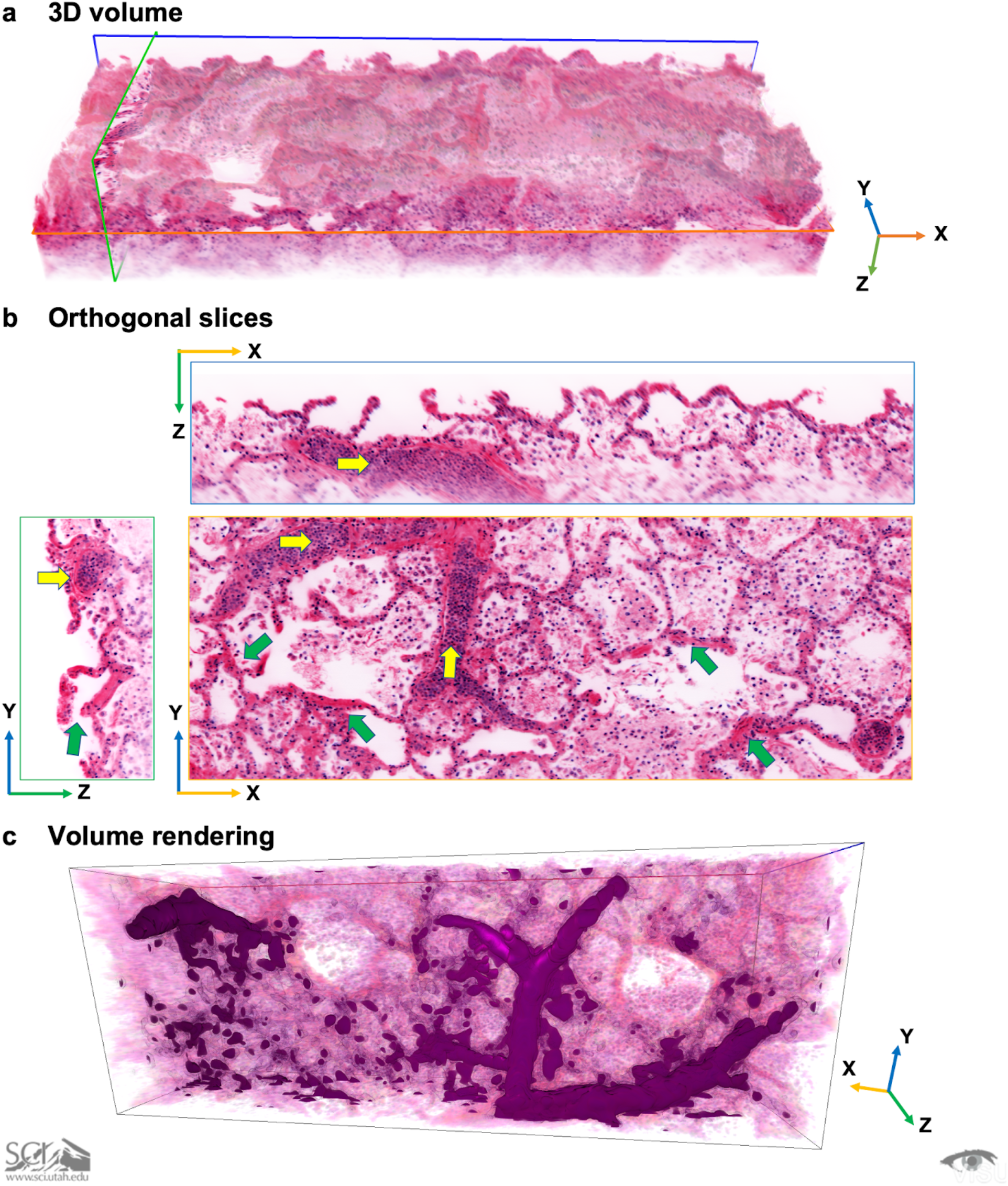
3D imaging reveals the extent of small pulmonary vessel microangiopathy. a) 3D volume of a 6 gigavoxel sub-volume of the tissue with selected orthogonal slices. The natural histologic surface topography of the sample is apparent. b) Orthogonal slices (XY, XZ, YZ) corresponding to the color-coded slices shown in the volume of (a). Platelet-rich clots with adherent mononuclear cells are observed in a medium-sized vessel surrounding a large alveolus (yellow arrows). The extent of the clot is apparent in the different orthogonal views. Extensive fibrin-rich clots in the smaller capillaries surrounding the alveoli are also apparent (green arrows). c) Volume render of the tissue adjusted to highlight the extent of the highly cellular clot, demonstrating the size and extent of the clot in this small area of the larger imaged volume.

## Conclusions

In this work, we present 3-dimensional virtual histological imaging of a single gross sample of lung tissue taken from a COVID-19 victim at the time of autopsy. This sample is from one of an initial series of African American patients reported from University Medical Center, New Orleans.^3^ That report was notable for its findings of small vessel thrombi and microangiopathy with hemorrhage significantly contributing to death. The purpose of this work was to leverage tissue clearing and high-throughput light sheet microscopy, combined with fluorescent H&E-analog stains and pseudocoloring, to provide additional insights in support of earlier findings.

The value of 3D imaging in this context is that it allows for much finer sampling of the tissue compared to traditional 2D slide-based histology, which may provide additional information or context that can support standard histology. In our case, by being able to digitally search and select any arbitrary section from mm^3^ volumes of the tissue, and to render volumes of interest in 3D, we were able to confirm findings that may have been difficult on single 2D sections due to unlucky sampling, and to visualize the true extent of cytopathic viral changes and damage to the lung from COVID-19. Notably, we were able to confirm the finding of activated mature megakaryocytes in the small vessels of the lung by inspection of their large, multiple, lobular nuclei conforming to the shape of the vessel in 3 dimensions. Additionally, 3D inspection of cells that appeared to be large multinucleated cells on 2D slices, enabled more accurate determination of a cell cluster. Three-dimensional imaging also revealed the massive extent of small vessel thrombosis in the lung. While small deposits of fibrin may be associated with diffuse alveolar damage more generally, 3D imaging allowed us to visualize the pathologic extent of fibrin deposition and clot formation with inflammatory cell attachment, which was far more extensive than revealed in 2D alone, providing additional strong support for the independent role of pulmonary microangiopathy in the pathogenesis of COVID-19 apart from diffuse alveolar damage.

Taken together, these findings support the role of 3-dimensional imaging of cleared tissues for the study of COVID-19 histopathology. The technological approach used here follows that described previously by our group and others.^4–9,11,13^ The specific fluorescence contrast scheme used here to replicate H&E,^7^ like similar schemes reported previously,^4–6^ has been shown repeatedly to provide reliable and accurate analogs to traditional H&E contrast in fluorescence microscopy. While our images were collected with a specific variant of light sheet microscopy, similar images could be obtained at varying levels of throughput with other optical sectioning microscopes. This approach will not likely soon replace traditional 2D histopathology due to its relative complexity and the size of the data generated - for a single sample from a single patient, we generated images nearing 1 teravoxel in size and exceeding 1.5 TB of processed data for the sample. However, it could serve as a useful adjunct to traditional histopathology, as shown here. In particular, as the figures and supplemental videos show, inspection of the 3D data provides a more comprehensive and intuitive understanding of the changes due to disease. While this report is limited to a single sample from a single patient, the utility of the findings support continued study on tissues from multiple organs from a wider cohort of patients, and with the addition of specific molecular markers, which is the subject of ongoing work.

## Materials and methods

### Sample acquisition

De-identified formalin-fixed gross lung tissue blocks were obtained from autopsies performed at the LSU Health Sciences Center – University Medical Center. Consent for autopsy was non-restricted by the next of kin, and these studies on autopsy tissues were determined to be exempt by the IRB at LSU Health Sciences Center and Tulane University. All experiments were conducted in accordance with SARS-CoV-2 (Risk Factor 3)-specific protocols approved by the Tulane University Institutional Biosafety Committee. Samples were fixed in formalin for over 72 hours prior to sample preparation according to institutional biosafety protocols.

### Sample preparation for three-dimensional (3D) imaging

After fixation the sample was rinsed overnight in PBST solution made from phosphate-buffered saline (PBS, Sigma) with 0.2% Triton-X 100 (Sigma), and then stained and cleared following the protocol of Reder et al.^7^ Briefly, the sample was immersed in a 1:1000 v/v solution of TO-PRO 3 iodide (T3605, Thermo Scientific) in PBST overnight with light shaking. Afterwards, the sample was dehydrated in graded solutions of 0% (deionized water), 25%, 50%, 75% and 100% ethanol solution with deionized water (1 hour each), followed by a final 1 hour wash in fresh 100% ethanol. The dehydrated sample was stained with alcoholic Eosin solution (Surgipath 3801615, Leica Biosystems) diluted in 1:1,000 v/v ratio with 100% ethanol overnight, then rinsed in ethyl cinnamate (ECi, Sigma) twice for 1 hour each time to undergo tissue clearing and refractive index matching.^9^ Finally, the sample was incubated in fresh ECi and degassed under vacuum for 1 hour.

### Three-dimensional (3D) imaging with dual-view inverted selective plane microscopy (diSPIM)

The system used to acquire 3D images in this work was a customized dual-view inverted selective plane microscopy (diSPIM) system, originally developed by Hari Shroff and co-workers and commercialized by Applied Scientific Instrumentation (ASI, Eugene, OR).^12^ Our system has been modified for large cleared tissue imaging with adjustable illumination tube lenses and dual Cleared Tissue Objectives (CTO, ASI/Special Optics, 15.3X - 17.9X, NA: 0.37-0.43 over RI range of 1.33 to 1.55, working distance: 12 mm) in order to adapt to diverse immersion mounting media, as described previously.^8^ The system is fitted with a 488-nm laser source (Omicron, PhoxX; maximum power = 200 mW) for fluorescence excitation of Eosin and a 647-nm laser source (Omicron, PhoxX; maximum power = 140 mW) for fluorescence excitation of TO-PRO-3 iodide. During imaging, the lung sample was immersed in ECi in a custom sample chamber, and imaged from above using the standard diSPIM arrangement.^8,11^ No additional material or interface was introduced between the objective lens surfaces and the cleared tissue other than the ECi immersion media, and no compression was applied, enabling reconstruction of the natural tissue surface topography. Although the system can be used to acquire 2 orthogonal views of the sample for dual-view imaging and reconstruction,^8,12,14^ in this work only one view was acquired. Captured light-sheet images from a scientific CMOS camera (Hamamatsu ORCA-Flash 4.0v2) were transferred to a controller workstation (Dell T3610) and stored in an 8TB SSD array set as RAID 0 mode. The raw data comprised a stack of 221,400 individual images with a single frame size of 2,048 pixels by 2,048 pixels, giving a total raw volume of 929 gigavoxels with a total file size of 1.69 TB.

### Image Processing

The raw data acquired from the diSPIM system were transferred to a processing workstation (Dell T7910 with Xeon E5-2699 v3 processor and 128 GB RAM). The first step was to conduct affine transformation and interpolation to shear vertical light-sheet images by 45 degrees horizontally plane-by-plane with assistance of data batching and parallel computation in MATLAB.^15^ Afterwards, illumination correction was conducted along the depth of each image, and all stacks of reconstructed volumes were stitched together to form the whole volume in FIJI.^16^ Then, the two 3D volumes of fluorescence images corresponding to the eosin and TO-PRO-3 channels were combined into a pseudo-color image in order to mimic the color palette of hematoxylin and eosin using the Virtual Transillumination Microscopy approach developed by Giacomelli et al.^5^ Visualizations of the 3D images were performed on both the single channel as well as pseudo-H&E volumes. Full volume visualizations were realized using Amira (Thermo Fisher Scientific) on 4 times downsampled images. After regions of interest were identified, sub-volumes were loaded at full resolution into Amira and the OpenViSUS^17^ interactive framework for further visualization. Volume renders for Figure 2c were created with a mixture of the single channel nuclear (TO-PRO-3) and pseudo-H&E volumes to visualize the cellular and tissue architectural detail using standard texture-based volume rendering.^18^ The visualizations for hyaline-fibrin aggregate and infected pneumocytes used the pseudo-H&E volume with the volume rendering transfer function designed to give semi-transparent stroma. The transfer function of the nuclear field was designed to give high intensity values an opaque, purple color while making mid to low intensity values transparency. For the megakaryocyte and infected cell cluster visualization, the single channel nuclear volume was added with standard volume shading applied to give perceptual hints of nuclei shape. Rendering order was enforced to make the nuclei field appear on top, avoiding nuclei color attenuation from the stroma in the pseudo-H&E volume. Rendering of the small vessel pulmonary microangiopathy was performed in a two-pass approach. First the single channel nuclear volume was thresholded to give all mid-to-high intensity nuclei maximum intensity. After this thresholding, this new field was blurred (Gaussian with a 30 pixel support). This blurring removed the contribution of regions with low concentrations of nuclei while maintaining the regions with large clusters to isolate the high cell density in the vessel thrombus. A similar transfer function as above was applied to this new field.

## Acknowledgements

We would first and foremost like to acknowledge the patients who succumbed to this deadly disease, and thank their families for allowing these studies to go forth to benefit the good of all. We would also like to thank the OpenViSUS team, especially Giorgio Scorzelli and Dr. Valerio Pascucci, for their help in accelerating our visualization efforts. J.Q.B, B.S., and C.W. acknowledge financial support from the National Science Foundation and National Institutes of Health under grant NSF DMS-1664848. B.S. acknowledges financial support from the National Science Foundation under IIS 1657020.

