## Supplemental Information for "Multiscale 3-dimensional pathology findings of COVID-19 diseased lung using high-resolution cleared tissue microscopy"

### Supplemental Methods

**Videos.** Videos of the reconstructed volumes in Supplemental Video 1 and Supplemental Video 3 were generated using Amira (Thermo-Fisher Scientific) and recorded with ScreenToGif. Supplemental Video 2 was captured in the OpenViSUS framework by rotating each subvolume about the z-axis.

### Supplemental Figures

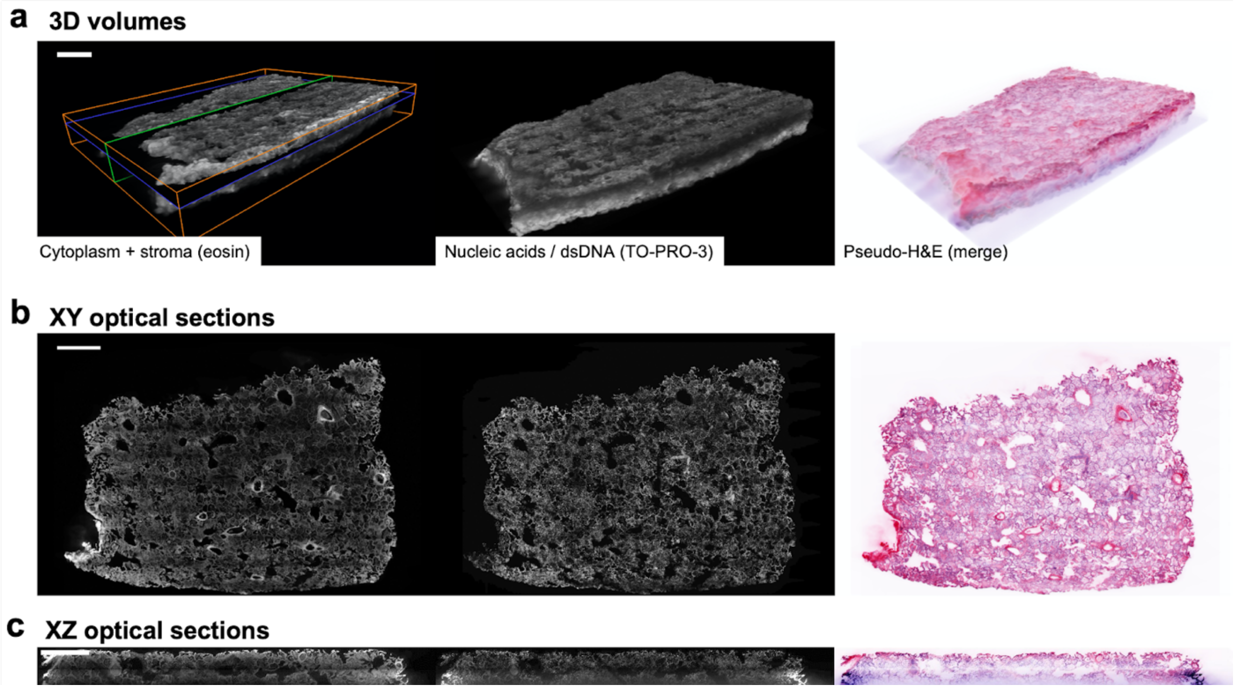


**Supplemental Figure 1. Generation of pseudo-H&E volumes (a) and sections (b, c) from individual acquisition channels of the entire volume corresponding to cytoplasm and stroma (eosin fluorescence) and nucleic acids (TO-PRO-3 stain, which is sensitive for dsDNA).**

**Supplemental Video 1.** Video of the subvolume from Figure 1b, demonstrating all orthogonal slices in each direction.

**Supplemental Video 2.** Video demonstrating lobular, multi-nucleated megakaryocyte in 3D.

**Supplemental Video 3.** Video of the subvolume from Figure 2a, demonstrating all orthogonal slices in each direction.
